# Simultaneous multicolor fluorescence imaging using duplication-based PSF splitting

**DOI:** 10.1101/2022.10.04.510770

**Authors:** Robin Van den Eynde, Fabian Hertel, Sergey Abakumov, Bartosz Krajnik, Siewert Hugelier, Alexander Auer, Joschka Hellmeier, Thomas Schlichthaerle, Rachel M. Grattan, Diane S. Lidke, Ralf Jungmann, Marcel Leutenegger, Wim Vandenberg, Peter Dedecker

## Abstract

We present a way to encode more information in fluorescence imaging by splitting the emission into copies of the original point-spread function (PSF), which offers broadband operation and compatibility with other PSF engineering modalities and existing analysis tools. We demonstrate the approach using the ‘Circulator’, an add-on that encodes the fluorophore emission band into the PSF, enabling simultaneous multicolor super-resolution and single-molecule microscopy using essentially the full field of view.

---

Fluorescence microscopy is a key technique in the life and materials sciences. A crucial challenge resides in maximizing the information content of the images. Several approaches have been developed to increase this content, including point-spread function (PSF) engineering, which encodes information in the pattern with which fluorophores appear on the detector. In optically-sparse samples, PSF engineering is highly attractive because it does not reduce the field of view or the temporal resolution of imaging. Commonly-used examples increase the axial resolution of the imaging via the introduction of astigmatism or more elaborate modifications [1, 2, 3, 4, 5], provide simultaneous dual-color imaging using complex phase shaping elements [6], or determine the emission spectrum of individual dyes at the cost of a reduction in field of view [7]. Recent work has also shown that the colors of individual emitters can be determined directly by leveraging the optical imperfections of the instrument [8], though with limited accuracy and the requirement for advanced analysis algorithms that must be trained and validated for every microscope and dye combination.

Several challenges complicate the general use of PSF engineering. Implementations in which the PSF can adopt a continuum of shapes require extensive data analysis that must accurately discern among many similar shapes (Figure 1a). The instrumentation may require either adaptive optics such as spatial light modulators, which come with their own limitations, or custom optical elements that must be carefully designed and manufactured for specific wavelength ranges and instruments. This also complicates the combination of multiple PSF-engineering techniques to encode multiple types of information at once. Overall, these issues fundamentally arise from the need to modify the PSF shape itself and limit the achievable information content. Avoiding direct shape manipulation would eliminate these restrictions, though this intuitively appears to be incompatible with the entire concept of PSF engineering.

**Figure 1:**
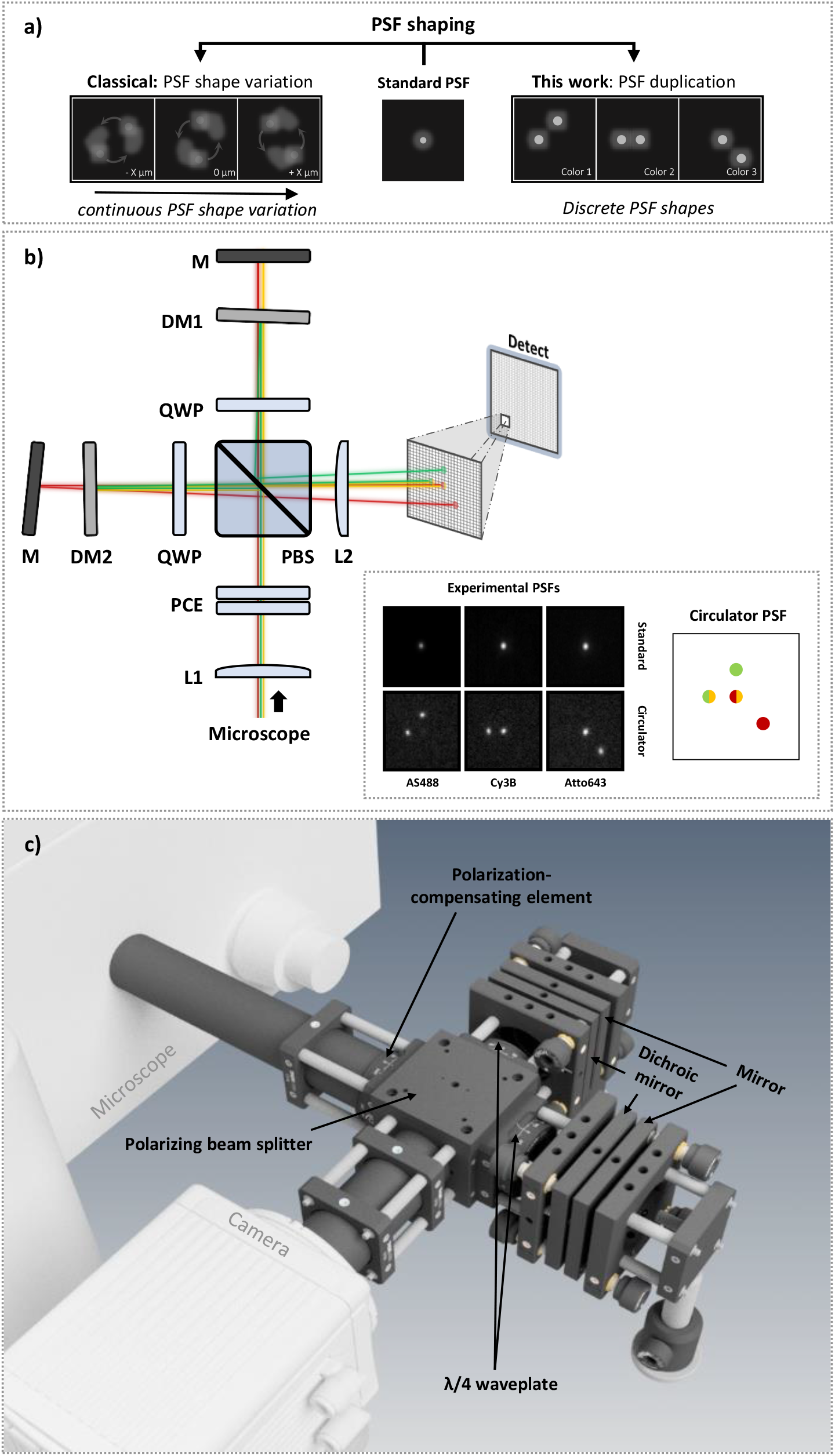
(a) Schematic illustration showing our duplication-based PSF engineering strategy and a more classical strategy based on the continuous variation of the PSF shape. (b) Optical layout of the Circulator and its components. The smaller panel shows a comparison between the unmodified PSF shapes and the corresponding Circulator PSFs for different fluorophores. (c) CAD rendering of the Circulator prototype used in this study.

We reasoned that these limitations can be avoided if, instead of trying to modify the shape of the PSF, we split the emission over two or more copies of the original PSF, each located at different positions on the detector. The desired information can then be encoded in the number, relative orientation, and/or intensities of these copies (Figure 1a). Such an approach poses several advantages when compared to alternative methods for PSF engineering: (i) it allows spectrally broadband operation because duplicating PSFs preserves the shape of the wavefronts and also does not reduce the sharpness of the original PSF, (ii) it does not interfere with other types of PSF engineering, (iii) it does not require new image analysis tools, and (iv) the theoretical localization accuracy of such a PSF is not reduced compared to that of the unmodified original PSF (Supplementary Note 1). By encoding this information into the relative orientation of two PSF lobes, our approach shares some similarity to the pioneering double-helix PSF engineering methodology [3].

We decided to explore this concept via the construction of a novel optical device that we call the ‘Circulator’ (Figure 1b and c). The Circulator uses PSF duplication to encode the emission color of individual fluorophores, by converting the native PSF, usually but not necessarily consisting of a single focused emission spot, into a pattern that consists of up to four different copies located at precisely-known distances and orientations relative to the position of the fluorophore. Our choice to encode the emission color reflects the profound importance of multicolor imaging. In conventional imaging, multicolor imaging is commonly achieved by sequential acquisition of the different color bands, which reduces throughput if the individual acquisitions take considerable time, as is the case for methodologies such as single-molecule localization microscopy (SMLM). Alternatively, dichroic image splitters can image distinct spectral bands onto distinct sensor areas of a camera. This works well, but results in a significant reduction of the field of view and registration overhead when using a single sensor, or can get tedious and costly when choosing a multi-camera approach in terms of mechanical instability, alignment requirements, and synchronization overhead, especially when imaging at higher spatial resolutions [9, 10]. Yet another strategy is to use temporal modulation of the light sources [11], which leads to a measurement slowdown since the fluorescence modulation of each emitter must now be captured over multiple acquired images, or the introduction of a diffractive or dispersive optical element [7], which requires bright emitters and reduces the field of view. Our approach, in contrast, does not suffer from these limitations, though it does share some of the inherent limitations of PSF engineering, such as a reduced ease of visual interpretation of the images, incompatibility with non-sparse imaging, and an increased sensitivity to background due to the increased spatial extent of the PSF.

The overall optical layout of the Circulator is shown in Figure 1b and c. The emission light coming from the microscope is collimated prior to propagating towards a polarizing beam splitter (PBS). A polarization-compensating element (PCE) in front of the PBS can be added to compensate for residual polarization and achieve an equal splitting of the incident light between the two optical arms (Supplementary Note 2). In each of these optical arms, the emission passes through a quarter-wave plate (QWP) located at a 45° orientation and is reflected by either a dichroic mirror or a subsequent broadband mirror depending on its wavelength, before passing the QWP and PBS again. The double pass through the QWP rotates the polarization of the light by 90° and ensures that it is transmitted/reflected by the PBS as appropriate and imaged onto the camera. The net result is that the imaging PSF consists of up to four different precisely-positioned copies of the original PSF, two defined by the dichroic mirrors and two more by the broadband mirrors. The relative intensities of each of these copies depends on the wavelength of the emission and the reflectivity of the dichroic mirrors, allowing the emitter color to be determined by straightforward visual or algorithmic inspection. The inset in Figure 1b shows measured PSFs for different dyes, in which one of the dichroic mirrors reflects green and orange light, and the other reflects only green light (Supplementary Figure S1). A detailed optical characterization of the device revealed that about 90% of the fluorescence is transmitted and that essentially no optical distortions are introduced (Supplementary Note 3). This basic concept shares some similarity with prior approaches that create additional emission spots via the introduction of diffractive optical elements [12, 13], though our approach preserves the PSF shape and is considerably more light-efficient.

We applied the Circulator to simultaneous three-color SMLM using DNA-PAINT. We identified Abberior Star 488, Cy3B, and Atto643 as suitable dyes for the green, orange, and red emission channels, respectively. Figure 2 shows an example three-color DNA-PAINT image obtained on a COS-7 cell in which the microtubules, vimentin, and clathrin were stained. A movie of the recorded data is available as Supplementary Information Movie 1. Since the observed PSFs consist of copies of the input PSF, the resulting images can either be analyzed using existing algorithms that detect the native PSF, and then post-processed using straightforward software that connects the correct relative lobe positions, or using a more dedicated analysis that directly recovers the different PSF patterns (Supplementary Note 4). Using this approach, we were able to acquire three color channels simultaneously, speeding up the overall imaging acquisition by a factor of three while retaining essentially the full field of view.

**Figure 2:**
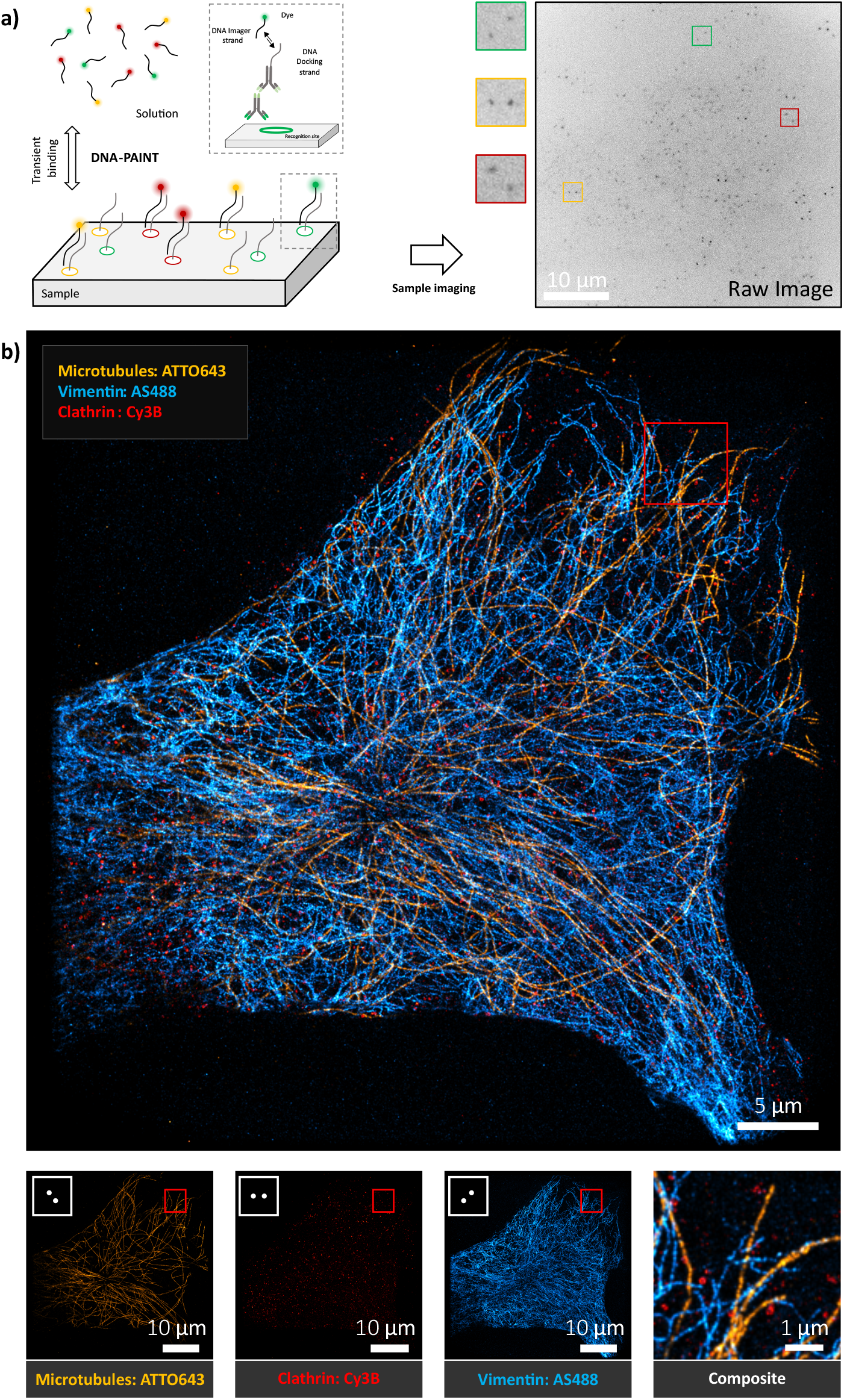
(a) Schematic illustration of DNA-PAINT imaging including an example fluorescence image in which three emitters of different colors are highlighted. (b) Top: Three color image of a DNA-PAINT Sample acquired using the Circulator. Bottom: Left to right, single-color channels showing microtubules, clathrin and vimentin structuring and a composite visualization. Microtubules, vimentin and clathrin were labeled with ATTO643, Abberior Star 488 and Cy3B respectively. Image acquisition time: 84 minutes.

A possible disadvantage is that the increased number of PSF spots reduces the maximal emitter density allowed before the analysis accuracy is reduced. However, even in a naive implementation the information content and throughput are still increased, because the number of spots is doubled but three different dyes can be distinguished. Furthermore, the Circulator PSFs are exactly known since they are determined by the mechanical alignment of the instrument, and as a result there is latent information in the image that can be used to resolve individual emitters even from dense regions. We analyzed this effect using an analysis based on machine learning (Supplementary Note 5), finding that the extra emission lobes can indeed lead to a lower analysis performance though to a much lower degree than could be naively expected.

We also combined our Circulator with superresolution optical fluctuation imaging (SOFI) [14, 15] in order to deliver multicolor high-resolution images at much higher emitter densities. SOFI relies on the statistical analysis of multiple images acquired from a sample labeled with emitters that show fluorescence dynamics, including DNA-PAINT samples [16]. Importantly, the SOFI analysis can also be extended to selectively extract emitters with a particular PSF shape by appropriate selection of the cross-cumulants used in the calculation [17] (Figure 3a and Supplementary Note 4.3). We applied this approach to DNA-PAINT samples exposed to high concentrations of the fluorophores. A movie of the raw data is provided as Supplementary Information Movie 2. An average image of the acquired images readily shows the image duplications created by the PSF copies (Figure 3b). From this single dataset, the SOFI analysis directly retrieves the three different high-resolution images showing the distinct emitters (Figure 3c). While the resolution of the SOFI images is lower than that of SMLM, the high emitter densities of this data made it possible to acquire these images approximately 50 times faster.

**Figure 3:**
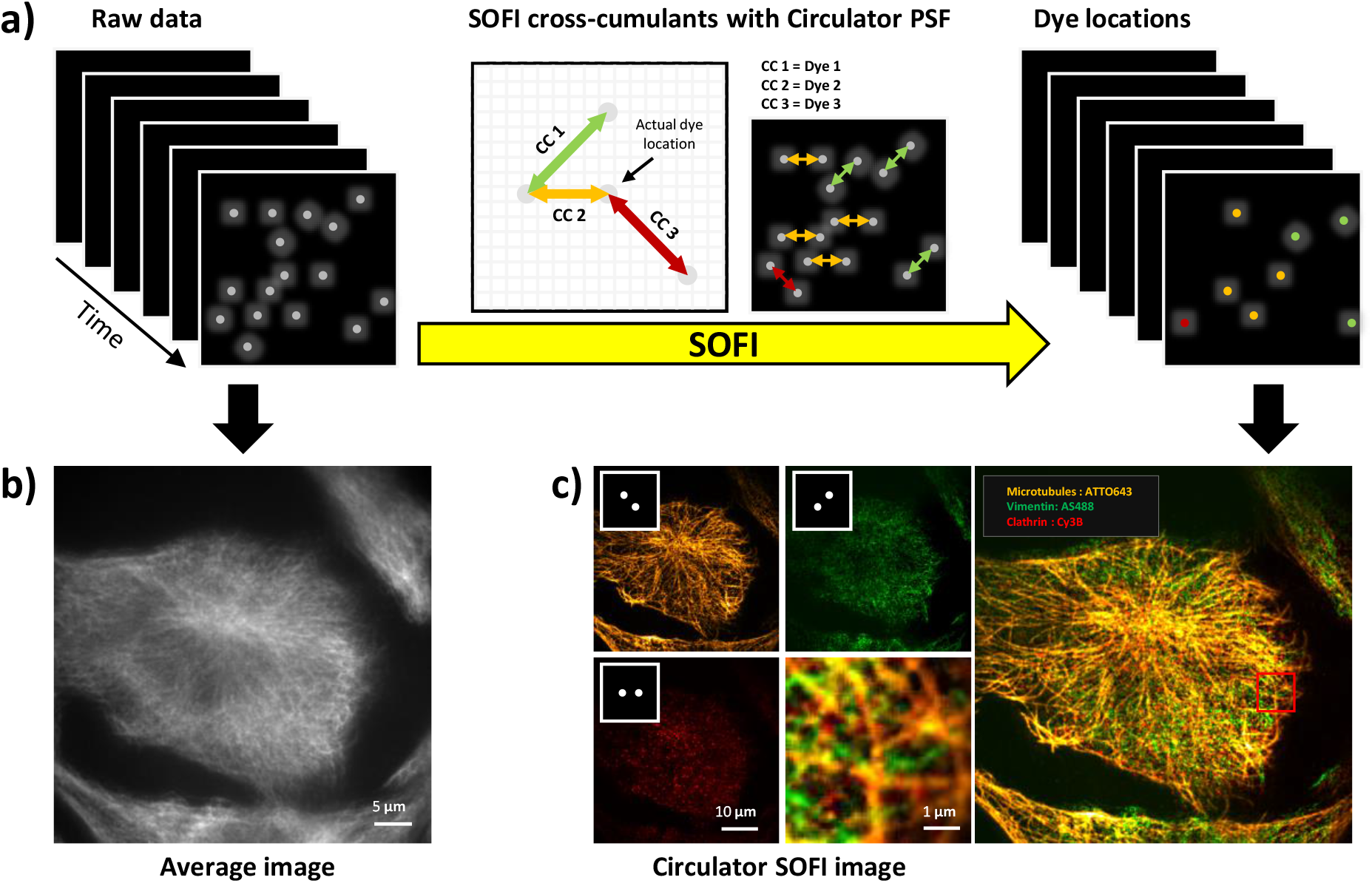
(a) Overall concept of SOFI-based imaging using the Circulator. (b) Fluorescence image obtained by calculating the average of all acquired fluorescence images. (c) SOFI images obtained from a single acquisition performed on cells in which microtubules, vimentin and clathrin were labeled with ATTO643, Abberior Star 488 and Cy3B respectively. Image acquisition time: 100 seconds.

We next performed single-particle tracking (SPT) experiments, which are typically performed on live or dynamic systems. Visualizing multiple components within a consistent state of the sample ideally requires simultaneous multicolor imaging, especially if interactions between these components are to be resolved. We evaluated the performance of our Circulator on an SPT model system consisting of a supported lipid bilayer on which cholesterol-anchored and fluorescently-labeled DNA origami molecules can freely diffuse (Figure 4a). Using this approach, we were able to monitor the different labeled constructs in parallel while straightforwardly discerning the color of each emitter based on their PSF shapes (Figure 4b and Supplementary Information Movie 3.

**Figure 4:**
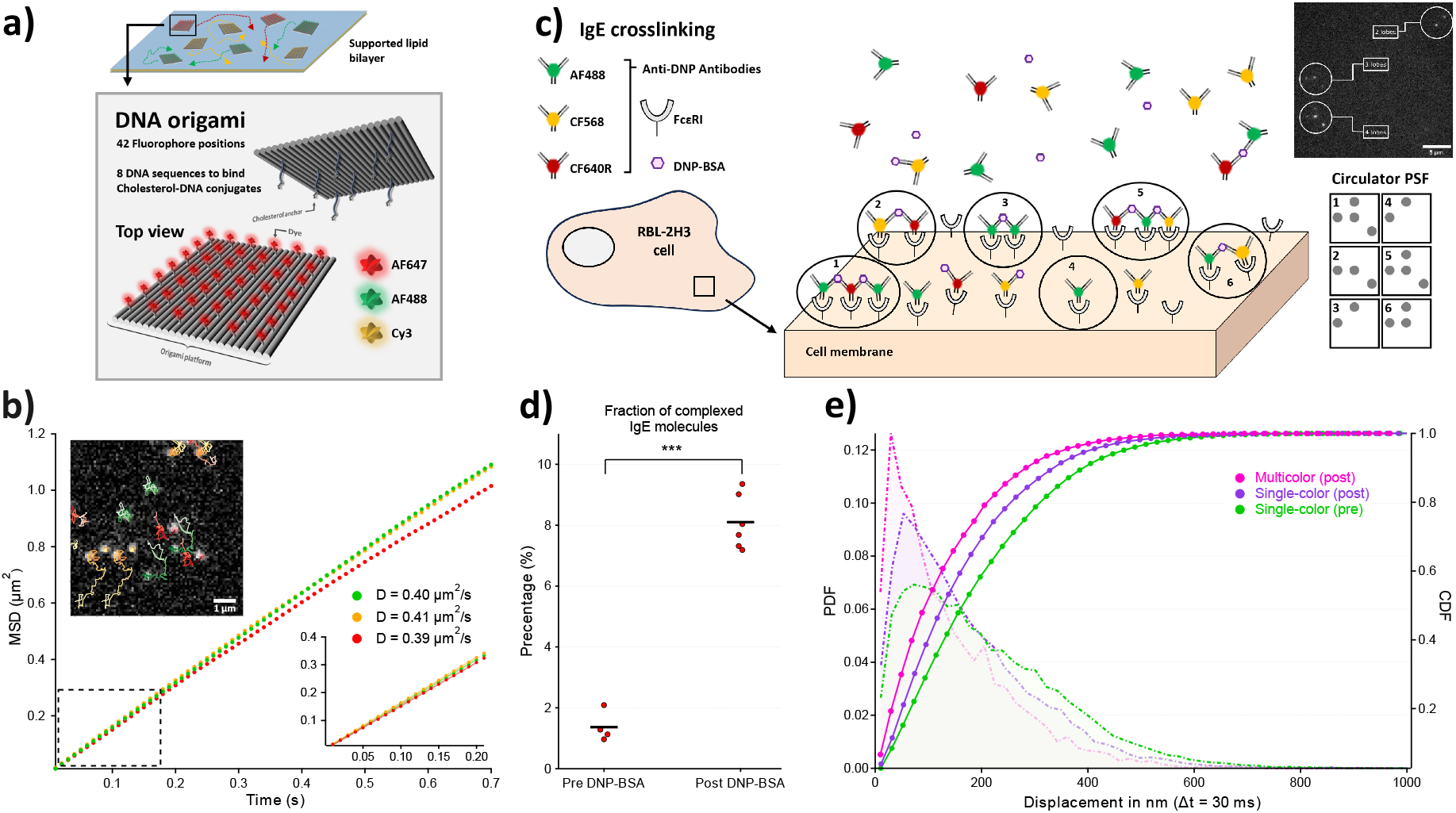
(a) A schematic description of DNA origami rafts on top of a lipid bilayer. The rafts are free to diffuse at the bilayer interface and are attached via cholesterol anchors. Each raft is stained with fluorophores of a single color. (b) Example SPT traces obtained for the freely diffusing origami rafts, as well as MSD plots for each color and the calculated diffusion coefficients. (c) Conceptual illustration of IgE crosslinking at the cellular membrane, the different PSF shapes this may create, as well as an example acquired fluorescence image. (d) Fraction of molecules that appear as double- or multilobe PSFs before- and after-stimulation with DNP-BSA. The unpaired t-test returned a *t*-value of 13.72, corresponding to a *p*-value of 7.6 ⋅10^−7^, showing a significant difference between two populations. (e) Step-size PDFs and CDFs for labeled IgE molecules pre- and post-addition of DNP-BSA. Multilobe fluorophores are not shown in the resting state due to insufficient statistics, which is consistent with a lack of aggregates before DNP-BSA addition.

We next turned our attention to SPT of a membrane receptor, FcεRI, on live cells. FcεRI is the primary immunoreceptor on mast cells responsible for the allergic response [18]. It binds antigen-specific IgE and is activated when multivalent antigen crosslinks multiple IgE-FcεRI into aggregates. We used the Circulator to observe IgE mobility and the formation of aggregates, by visualizing the IgE-FcεRI complex using anti-dinitrophenol (DNP) IgEs that were labeled with one of three different fluorophores (AF488, CF568 or CF640R). For these experiments, fluorescent IgE was added to RBL-2H3 cells at a concentration low enough to maintain a density suitable for single-molecule imaging. Individual IgEs could be readily visualized (Supplementary Information Movie 4), though the molecules were difficult to track for extended periods of time due to their low brightness and comparatively fast photodestruction. Nevertheless, we were able to investigate the association states of the observed particles by determining whether the observed fluorescence spots were consistent with single-color (two PSF lobes) or multicolor PSFs (three or four PSF lobes, indicating the presence of associated IgE molecules) (Figure 4c). We found that the particles were present primarily as single-color emitters in resting cells, though a strong increase was observed post DNP-BSA addition, consistent with receptor aggregation (Figure 4d). The ratio *r* of the number of particles appearing with four emission lobes to the number of particles appearing with three emission lobes was equal to 0.5 ± 0.19 (Supplementary Figure S2, indicating that the aggregates on average contain two IgE molecules (Supplementary Note 6). This also allowed us to calculate that approximately 8.1 ± 0.9% of the total amount of IgE molecules is present in such an aggregate.

These measurements also allowed us to calculate the diffusion dynamics for the monomeric and aggreggated receptors. Figure 4e shows the probability distribution functions (PDFs) and cumulative distribution functions (CDFs) of the displacements observed for individual single- and multicolor particles, displaying a clear reduction in mobility upon stimulation with DNP-BSA. This is consistent with previous SPT experiments showing that DNP-BSA addition causes receptor slow down [19], but here we are able to partially identify monomers from aggregates on the same cell. Furthermore, while some aggreggates will be misclassified by the analysis if they are, for example, labeled with two fluorophores of the same color, we estimate that the number of such misclassified particles is only 1.4% of the total number of observed particles. This is not sufficient to explain the observed decrease in single-color particle mobility evident from the CDF post-stimulation, indicating a reduction in IgE mobility that is not purely due to the formation of IgE homoaggreggates. We speculate that this reduced single-color FcεRI mobility may be due to changes in membrane compartmentalization in response to signaling by crosslinked receptors, but more work is needed to determine the underlying mechanism. Overall, these experiments show that our approach can readily increase the information content extracted from dynamic samples.

In conclusion, we have presented a new approach for PSF engineering that splits the emission light to create faithful copies of the native PSF. This new approach supports operation over a broad color spectrum, is compatible with other PSF engineering approaches, and does not require the development of new image analysis algorithms. We then demonstrated our approach by implementing the Circulator, an add-on module that encodes the emission color using only off-the-shelf components. The Circulator enabled simultaneous multicolor acquisitions using SMLM and SOFI, delivering a relative speedup up to the number of fluorophores used.

The device can also be adapted to image different dyes and processes by adjusting the dichroic mirrors that are used. In principle, the design could also be used to estimate the emission spectra of the emitters via a ratiometric analysis of the spot intensities. Beyond SMLM and SOFI, we expect the Circulator to be useful in any technique that involves sparse (single-molecule) emitters with different emission wavelengths, such as smFRET, single-particle tracking (SPT), and various in-situ ‘omics’ approaches, such as smFISH or MERFISH that can deliver sparseness via an intrinsically sparse spatial distribution of the labeled structures. Overall, we expect that the Circulator can readily increase the imaging information content and throughput of single-molecule or sparse imaging at very modest cost.

## Online Methods

### Hardware

The Circulator optics consisted of an optically-contacted polarizing beam splitter optimised for 400-700 nm (Bernhard Halle #PTW 25 OK), a quarter waveplate in each circulator arm positioned at 45° (Bernhard Halle #RAC 3.4.15), two mirrors (Newport #5101-VIS); a relay lens that collimates the light (Thorlabs #AC254-150-A); and an imaging lens (Thorlabs #AC254-060-A). The transmission/reflection spectra of the used dichroics are shown in Supplementary Figure S1. The mirrors were positioned near the common focal plane of the lenses. The polarization-compensating element (PCE) on our system consisted of a quarter-wave plate (Bernhard Halle #RAC 3.4.15) together with a half-wave plate (Thorlabs #AHWP05M-600). The device was coupled to an Olympus IX83 inverted microscope with a cellTIRF module equipped with 150 mW 488 nm, 150 mW 561 nm and 140 mW 640 nm lasers, an UAPON 150XOTIRF objective, manufacturer-installed quadband dichroic (Semrock #DI01-R405/488/561/635) and emission filter (Semrock #FF01-446/523/600/677), and a Hamamatsu Orca flash 4.0 V2 sCMOS camera. The projected pixel size of the camera was 108.3 nm.

### DNA-PAINT

The DNA-PAINT sample was purchased from Massive Photonics and contained human fibroblasts (InSCREENeX CIhuFIB). The antibody labeling was performed by Massive Photonics. Labeling of microtubules; primary antibody: anti-Tubulin alpha-Tubulin clone (Thermo Fisher Scientific #YL1/2) used in a dilution 1 in 200, secondary antibody: custom conjugated polyclonal anti-rat IgG (D1 docking strand). Labeling of clathrin; primary antibody: anti-Clathrin heavy chain (Abcam #ab21679) used in a dilution 1 in 400, secondary antibody: MASSIVE-AB 1-PLEX with anti-Rabbit IgG (D2 docking strand). Labeling of Vimentin; primary antibody: anti-Vimentin (Abcam #ab24525) used in a dilution 1 in 400, secondary antibody: custom conjugated polyclonal anti-chicken IgY (P7 docking strand). 90 nm gold fiducials for sample drift correction were provided by Massive Photonics and added following their instructions. The DNA imaging strands for Microtubule, Vimentin and Clathrin imaging were coupled to the ATTO643, Abberior Star 488 and Cy3B dyes respectively. ATTO643 and Cy3B dyes were provided by Massive Photonics. Abberior Star 488 was coupled to DNA by IBA. The imaging and docking strands used with Abberior Star 488 were “P7” as described in ref. [20]. SMLM raw data was obtained using a buffer containing: 200 µl buffer C (1X PBS pH 8, 500 mM NaCl, pH 8), 1.5 µl Imager 1-Atto643 (8 nM), 6 µl Imager 2-Cy3B (2 nM) and 6 µl P7-AS488 (200 nM). Imaging was performed using a total internal reflection (TIR) illumination. The SMLM analysis leading to Figure 2b was performed on 100.000 images with an exposure time of 50 ms per image. SOFI raw data was obtained using a buffer containing: 200 µl buffer C complemented with 8 mM KI (Thermo Scientific 373651000) and 300 mM MgCl_2_ (Product nr: 25108.295 - VWR), 2 μl 100× Protocatechuate 3,4-dioxygenase (PCD; Sigma-Aldrich #P8279-25UN), 5 μl 40× Protocatechuic acid (PCA; Sigma-Aldrich #37580-25G-F), 4 μl 50× Trolox (Sigma-Aldrich 238813-1G), 6 μl Imager 1-ATTO643 (50 μM), 12 μl Imager 2-Cy3B (50 μM) and 2 μl P7-Abberior Star 488 (5μM). 100x PCA and 40x PCA are prepared as described in ref. [20]. 50× Trolox was prepared by adding 100 mg of Trolox, 860 μl of methanol (Honeywell #24299-2,5L) and 690 μl of NaOH (1 M; Fisher Scientific #S/4880/60) in 6.4 ml of MilliQ water. Imaging was performed using a highly inclined and laminated optical sheet (HILO) illumination. The SOFI analysis shown in Figure 3c was performed on 2000 fluorescence images with an exposure time of 50 ms per image.

### Origami single-particle tracking

#### Assembly of DNA origami platforms

DNA origami structures (100×70 nm) were assembled in a single folding reaction carried out in a test tube (AB0620 - ThermoFisher Scientific) with 10 µl of folding mixture containing 10 nM M13mp18 scaffold DNA (Tilibit), 100 nM unmodified oligonucleotides, 500 nM fluorescently labeled oligonucleotides (AF488, Cy3, AF647 - Integrated DNA technologies), and folding buffer (5 mM Tris (ThermoFisher Scientific AM9855G), 50 mM NaCl (ThermoFisher Scientific AM9759), 1 mM EDTA (ThermoFisher Scientific AM9260G), 12.5 mM MgCl2) (ThermoFisher Scientific AM9530G)) [20]. At sites chosen for either fluorophore (42 positions spaced 10nm apart) or cholesterol anchor (8 positions spaced 20 nm apart) attachment, staple strands were elongated at their 3’-end with 21 or 25 bases, respectively. DNA origami were decorated with either of 3 different fluorophores (AF488, Cy3 or AF647). For this, 3× molar excess of respective fluorescently labeled oligonucleotides complementary to corresponding attachment sites on the DNA origami were added to the DNA folding mixture. DNA origami were annealed using a thermal protocol (90°C, 15 min ; 90°C - 4°C, 1°C/min ; 4°C, 6h) and purified using 100kDa Amicon Ultra centrifugal filters (Merck UFC510096). DNA origami were stored up to 4 weeks at -20°C.

#### Preparation of functionalized planar supported lipid bilayers (SLBs)

Vesicles containing 100% 1-palmitoyl-2-oleoyl-sn-glycero-3-phosphocholine (Avanti Polar Lipids #POPC) were prepared at a total lipid concentration of 0.5mg ml-1 as described [21] in 10x Dulbecco’s phosphate-buffered saline (PBS) (Sigma Aldrich #D1408-500ml). Glass coverslips (Menzel #1.5, 24×60 mm) were plasma cleaned for 5 min and attached Ibidi sticky-Slide 8-well chambers (Ibidi #80828). Coverslips were incubated with a fivefold diluted vesicle solution for 10 min, before they were extensively rinsed with PBS (Sigma Aldrich #D1408-500ML). For functionalization, SLBs were first incubated for 60 min with cholesterol-oligonucleotides (Integrated DNA technologies) at a final concentration of 0.1 µM complementary to the elongated staple strands at the bottom side of the DNA origami and then rinsed with PBS. DNA origami were incubated at a final concentration of 0.5 nM on SLBs for 60 min at ambient temperature.

#### IgE-FcεRI Colocalization

Approximately 20.000 RBL-2H3 cells were seeded into 8-well chambers (Cellvis #C8-1.5H-N), using supplemented MEM (Thermo Fisher Scientific #51200-038 supplemented with 10% Fetal Bovine Serum Thermo Fisher Scientific #10270106, 1% Penicillin-Streptomycin Thermo Fisher Scientific #15140122 and 2% L-Glutamine/GlutaMAX Thermo Fisher Scientific #35050038), one day prior to the experiment. The cells were washed three times with 200 μl of Hank’s Balanced Salt Solution (ThermoFisher Scientific #14025092, supplemented with BSA Sigma #A9647-100G to 0.05% w/v the day of imaging). Anti-DNP IgE [22] was conjugated to AF488, CF568, or CF640R fluorophores using NHS-ester chemistry [23]. The conjugated antibodies were diluted in HBSS (+BSA) and mixed to yield the following concentrations: 0.033 ng/ml AF488 IgE, 0.068 ng/ml CF568 IgE and 0.918 ng/ml CF640R IgE. Cells were labeled by adding 200 μl of this IgE premix to the well. Imaging of resting cells was performed for 5 to 15 minutes after IgE addition. Cells were then stimulated with the addition of 100 μl of 0.3 μg/ml DNP-BSA crosslinker (Thermo Fisher Scientific #A23019; stock solution: 200 μg/ml in PBS + 50 mM HEPES VWR #0511-1KG) to yield a final concentration of 0.1 μg/ml DNP-BSA. Imaging was performed for 2 - 15 minutes after DNP-BSA addition. Imaging was performed at room temperature.

### Data analysis

#### Emitter localization and color determination

The acquired fluorescence image sequences for SMLM reconstruction were filtered using a pixelwise sliding window median filter with a window size of 1,000 images, discarding the first 500 and last 500 images in the data for which the median cannot be calculated. This median filtering was applied to eliminate the contributions of fiducial markers and background emission. A uniform (flat) offset of 5,000 counts was then added to this filtered dataset in order to avoid the presence of negative values in the downstream postprocessing. The resulting image stack containing 99,000 images was then processed with the U-net based architecture described in Supplementary Note 4.2 and postprocessed using the ThunderSTORM SMLM analysis software [24]. The following parameters in ThunderSTORM were adjusted for appropriate emitter fitting: no noise removal was utilized, and the threshold for peak detection was set at 30 for each of the colors. The U-net was trained on data designed to resemble that observed in the SMLM experiments.

Sample drift in the SMLM acquisitions was corrected by separately analyzing the acquired (non-median-filtered) image stack using ThunderSTORM with the default settings to localize the bright fiducials. Localizations originating from fiducials were identified visually based on their brightness and persistence duration. Each fiducial furthermore four distinct localizations since it appeared with a four-lobe PSF, which were treated as separate particles. For every fiducial, displacements Δ*x* and Δ*y* relative to its position in its first frame of appearance were calculated for each image. We then calculated the overall shift of each image relative to the position of the first image, as the mean of all displacements of the fiducials localized within that frame. We then smoothed these shifts using a third-order Savitzky-Golay filter with a window size of 101 images so as to remove local jitter arising from the inaccuracy of emitter fitting. The resulting displacement estimates were then subtracted from the emitter positions localized in the corresponding images. For ease of visual interpretation, we removed the localizations originating from fiducials by removing all localizations arising within 1.5 pixels (162 nm) of their fitted positions.

The SMLM images were reconstructed using ThunderSTORM using the default settings and an eightfold adjusted magnification, and were rescaled to correct for unevenness in the illumination. Compound images showing all three color channels were then created by combining the different color channels in ImageJ.

#### SOFI analysis

The SOFI analysis was implemented using the Localizer software [25] in Igor Pro, using the built-in functionality for supplying specific cross-cumulant combinations. The individual cross-cumulants were selected by identifying the pixel combinations that would yield the highest expected signals for each channel, as described in Ref. [17].

#### SPT analysis

The SPT images were analyzed with the U-Net-based analysis as described in the SPT section and in Supplementary Note 4.2, yielding the positions of green, orange, and red emitters in the acquired images. No median filtering was used. The brightness of the emitters in the origami data was different compared to that observed in the SMLM data, which we compensated by first subtracting the mean value of each image and then multiplying these with a constant factor before applying the U-net-based analysis, where this factor was selected by optimizing the number of emitter molecules identified. The IgE data was not adjusted or rescaled, but was instead analyzed using a network specifically trained to recognize this data, as is likewise discussed in Supplementary Note 4.2.

The SPT analysis of the origami data was performed using custom Python code that assigned the positions detected in each image to previously detected particles using bipartite graph matching. The SPT analysis of the IgE data was performed using the Localizer software [25] in Igor Pro. To count as the same particle, emitters were allowed to disappear for up to 3 images (origami measurements) and 5 images (IgE measurements) in between two localizations, though no consecutive localizations could be more than 6 pixels (648 nm) apart for the origami data and 4 pixels (432 nm) for the IgE data. In the origami measurements, particles that had a displacement of less than 1 µm over their entire trajectory or that were observed in fewer than 16 images were removed from further consideration. For the IgE data, tracks obtained from particles observed in fewer than 3 images were removed from further consideration.

The origami dataset consisted of 948 green, 908 orange, and 755 red tracks. The mean-squared displacements (MSD) was computed by averaging the particle displacements (distances between consecutive localizations) calculated across the different tracks. The diffusion coefficient *D* of each species was determined by fitting the first 20 points in the obtained MSD curve with a linear fit, where the slope of this line was assumed to be equal to 4*Dt*.

In the IgE experiments, multilobe PSFs (emitter aggregates) were identified by identifying differently-colored trajectories that showed colocalization to within 1 pixel for at least four fluorescence images in which both particles could be detected. All such matching trajectories were combined into a single multicolor trajectory in which the particle location was set to the average of the locations of the tracks that could be observed in the corresponding fluorescence image. Data was acquired from 4 and 6 cells pre- and post-stimulation, respectively. In total, this yielded 2298 (2326) green, 2513 (2386) orange, 2755 (2639) red, and 41 (213) multicolor tracks pre- and (post-) stimulation.

The step size PDFs in the IgE data were calculated by calculating the distances the particles moved over in a particular time interval Δ*t* between the observations. These displacements were calculated for all tracks in their respective conditions (single vs. multicolor, and pre-vs. post-stimulation). We then calculated a weighted histogram of these displacements, where the weight associated with a given track was set to 1/*d*, where *d* is the number of displacements calculated for this track. This weighting assured that each particle had an equal contribution irrespective of the number of fluorescence images that it was observed in. The PDFs were then calculated by scaling the histograms such that the sum of all bins was equal to one. The CDFs were then calculated by creating a histogram in which the value of each bin was the sum of all the preceding bins in the PDF.

The number of IgE molecules in a complex (dimer or oligomer) and the average number of monomers in these complexes was calculated as detailed in Supplementary Note 6.

## Acknowledgments

We thank Arno Bouwens (KU Leuven) for fruitful discussions regarding the Circulator design and Toon Van Thillo (KU Leuven) for the preparation of alignment samples. We thank Yoav Shechtman and Elias Nehme (Technion) for discussions and insights regarding the use of machine learning for the data analysis. R.V.D.E. and S.H. thank the Research Foundation-Flanders (FWO Vlaanderen) for a doctoral and postdoctoral fellowship, respectively. We thank Derek Rinaldi for the generation of the fluorescent IgE. This work was supported by the European Research Council through grant 714688 NanoCellActivity, the Research Foundation-Flanders through grant G090819N, and the National Insitutes of Health via grant R35GM126934. B.K. thanks the Polish Ministry of Science and Higher Education’s Mobility Plus programme (1068/MOB/2013/0).

## Author contributions

W.V. and P.D. designed research. P.D. supervised research. M.L. invented the optical layout of the Circulator. B.K., R.V.D.E. and W.V. constructed the Circulator. R.V.D.E., F.H., and W.V. performed experiments and analyzed data with assistance from S.H.. W.V., S.A, and P.D. developed new analysis tools and algorithms and analyzed data. A.A., T.S., and J.H. created samples under the supervision of R.J. and performed initial experiments. D.S.L. and R.M.G. provided samples and interpretation for the SPT experiments. P.D. wrote the manuscript with input from W.V. and R.V.D.E.

## Data and Code Availability

Data and code are available upon reasonable request.

