## Supplementary information for "Simultaneous multicolor fluorescence imaging using duplication-based PSF splitting"

Robin Van den Eynde[1], Fabian Hertel[1], Sergey Abakumov[1], Bartosz Krajnik[1,2], Siewert Hugelier[1], Alexander Auer[3,4], Joschka Hellmeier[3,4], Thomas Schlichthaerle[3,4], Rachel M. Grattan[5], Diane S. Lidke[5], Ralf Jungmann[3,4], Marcel Leutenegger[6], Wim Vandenberg[1,*], Peter Dedecker[1,*]

[1] Department of Chemistry, KU Leuven, Belgium
[2] Department of Experimental Physics, Wroclaw University of Science and Technology, Wroclaw, Poland
[3] Max Planck Institute of Biochemistry, Martinsried, Germany
[4] Faculty of Physics and Center for Nanoscience, Ludwig Maximilian University, Munich, Germany
[5] Department of Pathology and Comprehensive Cancer Center, University of New Mexico, Albuquerque, United States of America
[6] Max Planck Institute of Biophysical Chemistry, Göttingen, Germany

#### Contents

**Supplementary figures** **2**

**1 Theoretical localisation precision of two-lobed PSFs** **3**

**2 The addition of a polarization-compensating element between the microscope body and Circulator** **5**

**3 Optical performance of the Circulator** **6**

**4 Analysis of Circulator-generated images** **7**

**5 Effect of the emitter density on the Circulator localization performance** **10**

**6 Calculation of the IgE aggregate sizes** **11**

### Supplementary figures

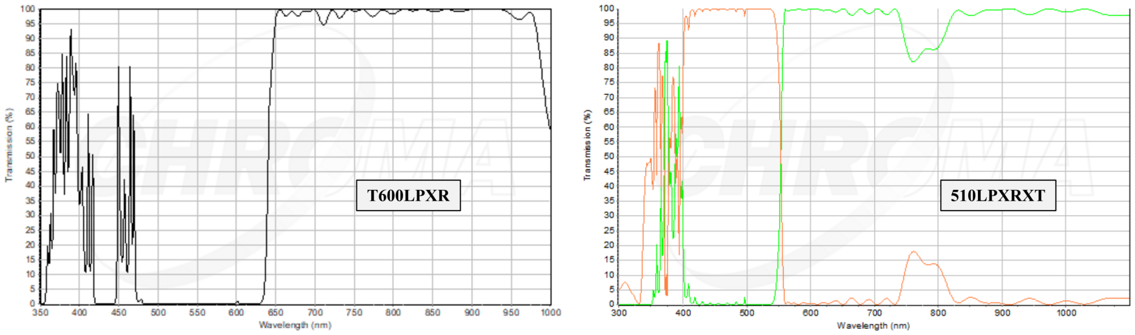

**Figure S1:** Transmission/reflection spectra of the dichroics used in the Circulator, at zero degree angle of incidence, as provided by Chroma. Left: Transmission of the T600LPXR dichroic. Right: Reflection (green) and transmission (orange) of the T510LPXRXT dichroic.

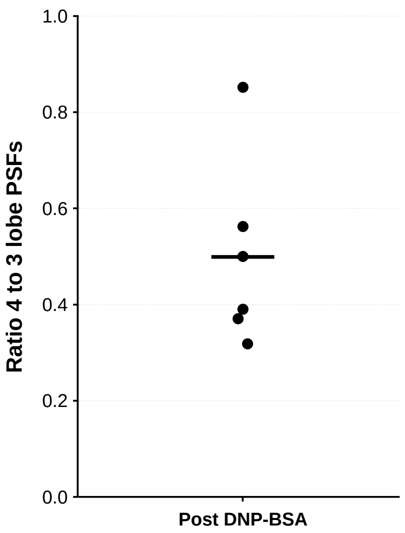

**Figure S2:** Ratio of the number of multicolor particles showing four emission lobes to the number of multicolor particles appearing with three lobes post-stimulation with DNP-BSA. Each marker represents an individual cell. The horizontal line shows the mean value of the data points.

#### Supplementary Note 1: Theoretical localisation precision of two-lobed PSFs

We sought to determine analytically how the theoretical localization precision of a single-lobe Gaussian PSF is changed when the PSF is converted into a two-lobe PSF, as is the case with our current implementation of the Circulator. We follow the procedure described in Ref. [1], where the limit of the $x$-localization accuracy is given by $\psi_x = \sqrt{[I^{-1}]_{11}}$ and the y-localization accuracy ($\psi_Y$) is given by $\psi_x = \sqrt{[I^{-1}]_{22}}$. In these expressions, $I$ is the Fisher information matrix, determined by:

$$I = N \begin{bmatrix} \int_{x,y} \frac{1}{\phi}\Big(\frac{d\phi}{dx}\Big)^2 & \int_{x,y} \frac{1}{\phi}\frac{d\phi}{dx}\frac{d\phi}{dy} \\ \int_{x,y} \frac{1}{\phi}\frac{d\phi}{dx}\frac{d\phi}{dy} & \int_{x,y} \frac{1}{\phi}\Big(\frac{d\phi}{dy}\Big)^2 \end{bmatrix} \tag{1}$$

where $N$ is the number of detected photons and $\phi$ the point spread function. We assume that the Circulator PSF can be described as the sum of two Gaussians with standard deviation $\sigma$, identical to the standard deviation of the non-Circulator PSF, that are furthermore separated along the $y$-axis by a distance $\delta$. More formally:

$$\phi = \phi_x \cdot [\phi_y^1 + \phi_y^2] \tag{2}$$

where $\phi_x$ and $\phi_y$ describe the PSF profiles along the $x$ and $y$ directions:

$$\phi_x = \frac{1}{\sigma\sqrt{2\pi}} e^{-\frac{1}{2}\frac{x^2}{\sigma^2}} \tag{3}$$

$$\phi_y^1 = \frac{1}{2\sigma\sqrt{2\pi}} e^{-\frac{1}{2}\frac{(y-\delta/2)^2}{\sigma^2}} \tag{4}$$

$$\phi_y^2 = \frac{1}{2\sigma\sqrt{2\pi}} e^{-\frac{1}{2}\frac{(y+\delta/2)^2}{\sigma^2}} \tag{5}$$

The additional factor 2 in the denominator of equations (4) and (5) reflects the fact that the light is split between both lobes. The derivatives of this PSF with respect to $x$ and $y$ are given by

$$\frac{d\phi}{dx} = -\frac{x\phi_x}{\sigma^2}[\phi_y^1 + \phi_y^2] \tag{6}$$

$$\frac{d\phi}{dy} = -\frac{\phi_x}{\sigma^2}\Big(y[\phi_y^1 + \phi_y^2] + \delta/2[\phi_y^1 - \phi_y^2]\Big) \tag{7}$$

We can now compute the elements of the Fisher information matrix by using the following well-known integrals:

$$\int_{-\infty}^{\infty} \phi_x dx = 1 \tag{8}$$

$$\int_{-\infty}^{\infty} x\phi_x dx = 0 \tag{9}$$

$$\int_{-\infty}^{\infty} x^2\phi_x dx = \sigma^2 \tag{10}$$

$$\int_{-\infty}^{\infty} \phi_y^1 dy = \int_{-\infty}^{\infty} \phi_y^2 dy = \frac{1}{2} \tag{11}$$

$$\int_{-\infty}^{\infty} y\phi_y^1 dy = \frac{\delta}{4} \tag{12}$$

$$\int_{-\infty}^{\infty} y\phi_y^2 dy = -\frac{\delta}{4} \tag{13}$$

$$\int_{-\infty}^{\infty} y^2\phi_y^1 dy = \int_{-\infty}^{\infty} y^2\phi_y^2 dy = \frac{\sigma^2}{2} + \frac{\delta^2}{8} \tag{14}$$

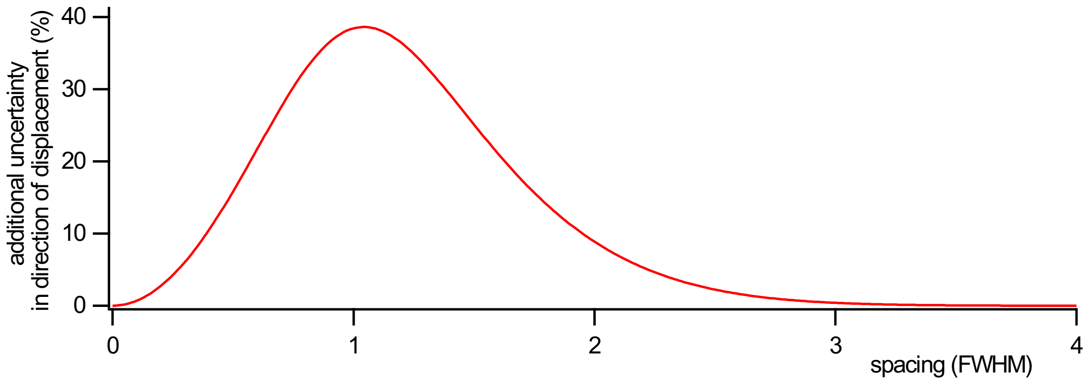

**Figure S3:** Graph of the additional uncertainty in the localization of a 2-lobe PSF in the direction of duplication in function of the sepparation between the copies (in units of the FHWM of the lobes)

leading to the following expressions:

$$\begin{aligned} I_{11} &= N \int_{x,y} \frac{1}{\phi}\Big(\frac{d\phi}{dx}\Big)^2 \\ &= \frac{N}{\sigma^4} \int_y \phi_y \int_x x^2 \phi_x \\ &= \frac{N}{\sigma^2} \end{aligned} \tag{15}$$

$$\begin{aligned} I_{12} = I_{21} &= N \int_{x,y} \frac{1}{\phi} \frac{d\phi}{dx} \frac{d\phi}{dy} \\ &= \frac{N}{\sigma^4} \int_x x\phi_x \int_y y[\phi_y^1 + \phi_y^2] + \delta/2[\phi_y^1 - \phi_y^2]) \\ &= 0 \end{aligned} \tag{16}$$

$$\begin{aligned} I_{22} &= N \int_{x,y} \frac{1}{\phi}\Big(\frac{d\phi}{dy}\Big)^2 \\ &= \frac{N}{\sigma^4} \int_x \phi_x \int_y \frac{(-y(\phi_y^1 + \phi_y^2) + \delta/2(\phi_y^1 - \phi_y^2))^2}{\phi_y^1 + \phi_y^2} \\ &= \frac{N}{\sigma^4} \int_y (\delta/2 - y)^2(\phi_y^1 + \phi_y^2) - 2\delta(\delta/2 - y)\phi_y^2 + \frac{(\delta\phi_y^2)^2}{\phi_y^1 + \phi_y^2} \\ &= \frac{N}{\sigma^4}\Big(\sigma^2 - \delta^2/2 + \delta^2 \int_y \frac{(\phi_y^2)^2}{\phi_y^1 + \phi_y^2}\Big) \end{aligned} \tag{17}$$

where eq. (17) was simplified by substituting $\Delta = \frac{\delta}{\sigma}$ and $v = y/\sigma$

$$I_{22} = \frac{N}{\sigma^2}\Big(1 - \Delta^2/2 + \Delta^2 \frac{1}{2\sqrt{2\pi}} \int_v \frac{e^{-(v+\Delta/2)^2}}{e^{-1/2(v+\Delta/2)^2} + e^{-1/2(v-\Delta/2)^2}}\Big) \tag{18}$$

Since $I$ is a diagonal matrix ($I_{12} = I_{21} = 0$), the inverse is given simply as $I_{11}^{-1} = 1/I_{11}$ and $I_{22}^{-1} = 1/I_{22}$. The limit on the localization precision will thus be given by

$$\psi_x = \frac{\sigma}{\sqrt{N}} \tag{19}$$

$$\psi_y = \frac{\sigma}{\sqrt{N}} G(\Delta) \tag{20}$$

here $\psi_x$ is the classical localization precision and $\psi_y$ is this same precision scaled with a function $G$ which depends solely on the separation between the lobes relative to the width of the lobes:

$$G = \left[1 - \frac{\Delta^2}{2} + \Delta^2 \frac{1}{2\sqrt{2\pi}} \int_v \frac{\exp -(v + \Delta/2)^2)}{\exp\left(-1/2(v + \Delta/2)^2\right) + \exp\left(-1/2(v - \Delta/2)^2\right)}\right]^{-\frac{1}{2}} \tag{21}$$

Figure S3 shows this additional localization uncertainty in the $y$ direction (the direction along which the two lobes are separated) as a function of the distance between the lobes, where this distance is expressed in units of the full width at half maximum (FWHM) of the lobes ($\approx 2.355\sigma$). When both lobes overlap perfectly ($\Delta = 0$), we find no additional uncertainty, as is to be expected. As the lobes start to separate, the localisation precision becomes worse compared to the single-lobe case, almost reaching a value $\sqrt{2}$ times higher at a separation of approximately 1 FWHM, before rapidly returning to the single-lobe fitting accuracy for further increasing distances. This result makes intuitive sense as during the intermediate (0 to 2 FWHM) regime the lobes overlap, making it difficult to unambiguously assign incoming photons to one of the two lobes. For larger distances, however, the theoretical accuracy in fitting a two-lobe PSF is exactly the same as that found for fitting an unmodified single-lobe PSF.

### Supplementary Note 2: The addition of a polarization-compensating element between the microscope body and Circulator

In principle, the unpolarized fluorescence emission is split in beams of equal intensity by the polarizing beamsplitter, leading to two emission spots that are equally bright. However, differing spot brightnesses can occur when the excitation light is linearly or elliptically polarized, when the fluorophores do not rotate freely, or when one of the polarization axes is preferentially transmitted or detected by the instrument. Furthermore, combining an emission polarizer with optical elements with a high numerical aperture, as is typical for the objectives used in high-resolution imaging, intrinsically introduces an ellipticity into the PSF [2]. To ameliorate such effects, we propose the insertion of a polarization-compensating element before the polarizing beam splitter ('PCE' in Main Text Figure 1). In principle, a broadband quarter waveplate should provide sufficient compensation possibilities to achieve uniform splitting. However, on the instrument used in this work, we found that an additional half-wave plate was necessary, most likely caused by a failure of the Olympus IX-83 body and optical components therein to fully maintain the polarization of the emitted light. In part, these originate from the differing S- and P-retardation of the light transmitted through the dichroic mirror that is present in the microscope (Figure S4). Both the quarter and half waveplates were rotated in order to empirically obtain symmetric PSF lobe intensities and shapes.

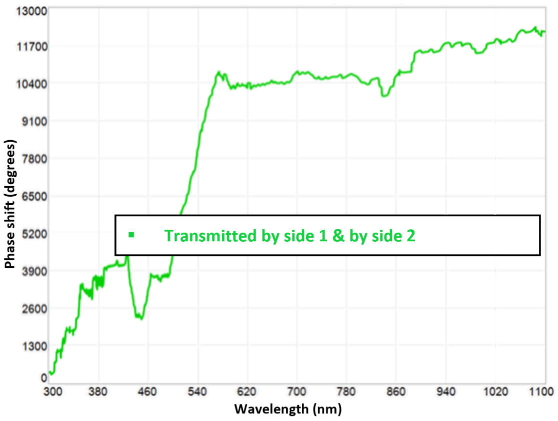

**Figure S4:** Phase difference between $p$ and $s$ light transmitted by the by the R405/488/561/635 dichroic in the used microscope. 'Transmitted by side 1 & by side 2' means that the light passes through both sides of the dichroic mirror, as is customary. (Figure courtesy Semrock).

### Supplementary Note 3: Optical performance of the Circulator

We performed a detailed characterization of the optical quality of the Circulator. First, we quantified the light loss, which is the amount of light that does not reach the detector as well as the amount of light that impinges on the detector at an unexpected position. To determine the first component, we removed the Circulator from the microscope and introduced a 532 nm laser (Thorlabs #CPS532) along the same path that light from the microscope would travel along (including the lenses denoted as L1 and L2 in Figure 1 in the main text). We quantified the total laser power using a power meter (Thorlabs #PM100D/S120-FC) before the Circulator and at the position where the camera would normally be, resulting in an 11.3% measured loss of light.

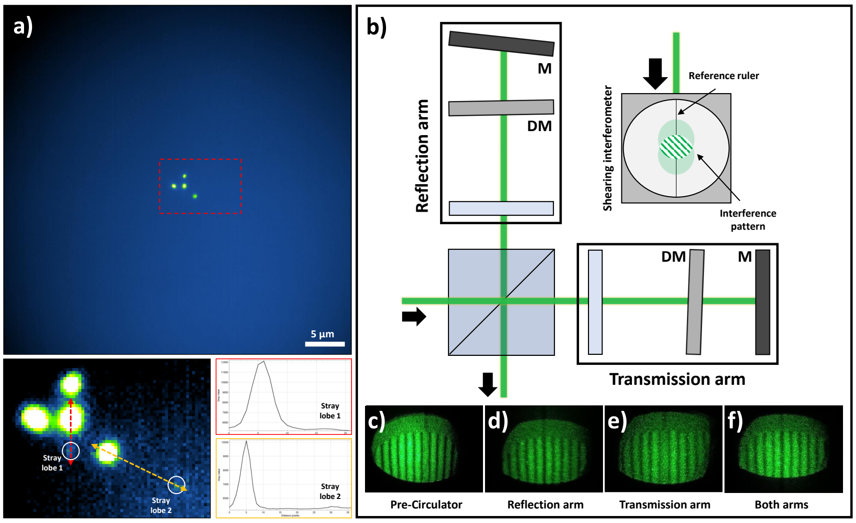

**Figure S5:** (a) Average of 50 fluorescence images acquired on a fluorescent bead emitting across all three spectral bands, using the Circulator and a per-image camera exposure time of 2 s. Two very low intensity stray lobes can be observed when the image is oversaturated (bottom). Line profiles show that the combined counts of these stray lobes is less than 1% of the total observed emission. (b) Optical characterisation of the Circulator using interferometry. A shearing interferometer, with ruled reference, is used to determine collimation and expose aberrations. All images were taken using a hand-held camera. (c) Pre-Circulator. Lineair lines that are parallel to the reference can be observed prior to the Circulator. These lines are straight, which indicates no apparent aberrations are present. (d–f) After passage through different parts of the device, the observed lines remain parallel to the reference and straight, showing that no or very little aberrations are introduced.

Next, we quantified the amount of light travelling in spurious directions due to stray reflections originating from intermediate surfaces, which should be observable as additional lobes in the PSF. We imaged isolated fluorescent beads (Thermo-Fisher 100 nm TetraSpeck Microspheres) illuminated using simultaneous 488, 561 and 651 nm light as seen in Figure S5a. Only the four Circulator lobes could be directly observed when performing a customary visualization of the images. By averaging over 50 consecutive images, we nevertheless uncovered two stray signals as highlighted in the oversaturated visualization and the accompanying line plots. Together these lobes constitute less than 1% of the total signal (calculated as the integrated intensity of the stray lobes divided by the integrated intensity of all expected lobes).

We also determined whether emission in a single color band could give rise to confusion by also appearing in lobes associated with different color bands. To verify this in practice, we spectrally isolated the single channels by illuminating the sample using a single laser and introducing a suitable emission filter in the microscope body (Chroma #et525/30m for 488 nm light, Olympus #ba575-625 filter for 561 nm light, and Semrock #FF01-446/523/600/677 filter for 640 nm excitation). We found that the lobes belonging to incorrect channels were so weak they could not be discerned from noise. Taking these three effects into account, we conclude that the total light loss of the Circulator is approximately 13%, which is in line with that expected from the optical design.

We then verified whether the Circulator introduced appreciable distortions of the optical wavefront. We did this using interferometry, leveraging a shearing interferometer (Thorlabs #SI050) to detect potential optical aberrations including defocus, spherical aberration, coma, and astigmatism [3]. We introduced a laser beam (Thorlabs #CPS532 expanded 2.5 times) as shown in Figure S5, imaged it onto a suitable shear-plate after the second pass (Thorlabs #SI100) and recorded the interference pattern, including the blocking of one of the 'arms' of the system. There was no observed difference between the light patterns observed before and after the Circulator (Figure S5c-f). Overall, we conclude that the Circulator does not introduce noticeable distortions into the imaging.

### Supplementary Note 4: Analysis of Circulator-generated images

The data generated by the Circulator require adapted or new analysis algorithms that can extract the information encoded in the generated PSFs. We propose three different ways in which this can be done: (i) adding post-processing routines to existing (non Circulator-aware) analysis algorithms, (ii) leveraging machine learning to generate synthetic single-color fluorescence images that can be analyzed using classical means, and (iii) PSF-aware SOFI analysis on high density data. We describe each of these approaches in the following sections.

#### 4.1 Analyzing Circulator data using post-processing of classical analysis results

The Circulator generates PSFs that contain two or more copies of the input PSF at well-defined relative positions, independent of the exact nature of this input PSF. Because the copies have the same shape as the unmodified PSF, though with a lower brightness, in principle any analysis software that is capable of detecting and analyzing the unmodified PSF is also capable of separately localizing the two copies belonging to a single Circulator PSF. Applying an SMLM analysis to data imaged with the Circulator, for example, will in principle result in a list containing two emitter localizations for each of the emitters in the sample. Given that the relative distances and orientations between the Circulator PSF copies are exactly known, this list can be post-processed using simple software that then yields both the actual emitter coordinates as well as the color of each of the emitters.

In a nutshell, this post-processing can be done in the following way. Let $\{\vec{x}_i\}$ be the set of PSF coordinates found by the analysis (twice as many coordinates as there are active fluorophores). For each such coordinate $\vec{x}$, verify whether the analysis also found another coordinate $\vec{x}$' at a position that would match the relative displacement associated with one of Circulator PSFs. If so, we remove $\vec{x}$ and $\vec{x}$' from the coordinate list and use our knowledge of the Circulator PSFs to add the actual emitter position and color to the fluorophore localizations. The procedure is then repeated until there are no more coordinates left or until no more matches are found.

Additional complexity can be added to this algorithm in order to increase its performance, such as requiring that the matched PSF coordinates also have the expected relative intensities, though the overall concept remains the same. Such a simple approach makes it comparatively easy to introduce the Circulator within an existing analysis pipeline, on the conditions that the emitter density is not too large to make the analysis ambiguous, and that the brightness of the emitters is sufficiently high so that the two PSF copies can be detected independently.

#### 4.2 Machine learning-based analysis

Machine learning based on convolutional neural networks (CNNs) has proven highly capable for the recognition and analysis of structural features within visually complex and/or noisy imaging data. They have also previously been applied to the analysis of high-density SMLM data (e.g. [4]), suggesting that they may function well also for the analysis of high-density Circulator measurements by learning the double-lobe nature of the PSFs. We elected to introduce the methodology in a hybrid approach in which the CNN takes an input fluorescence image measured using the Circulator and produces three new artificial fluorescence images that approximate what would be observed if each of the channels were observed separately with an unmodified (single-lobe) PSF (Figure S6a). These images can then be analyzed using established SMLM-analysis tools to yield the coordinates of the fluorophores.

We decided to introduce a CNN with the overall configuration shown in Figure S6b. It consists of a fully convolutional encoder-decoder network (FCN) that takes an input fluorescence image of size $N \times N$, but with a modified output that and produces three calculated images of the same size, separately showing the green, orange, and red emission channels. We also switched the activation within the convolutional layers

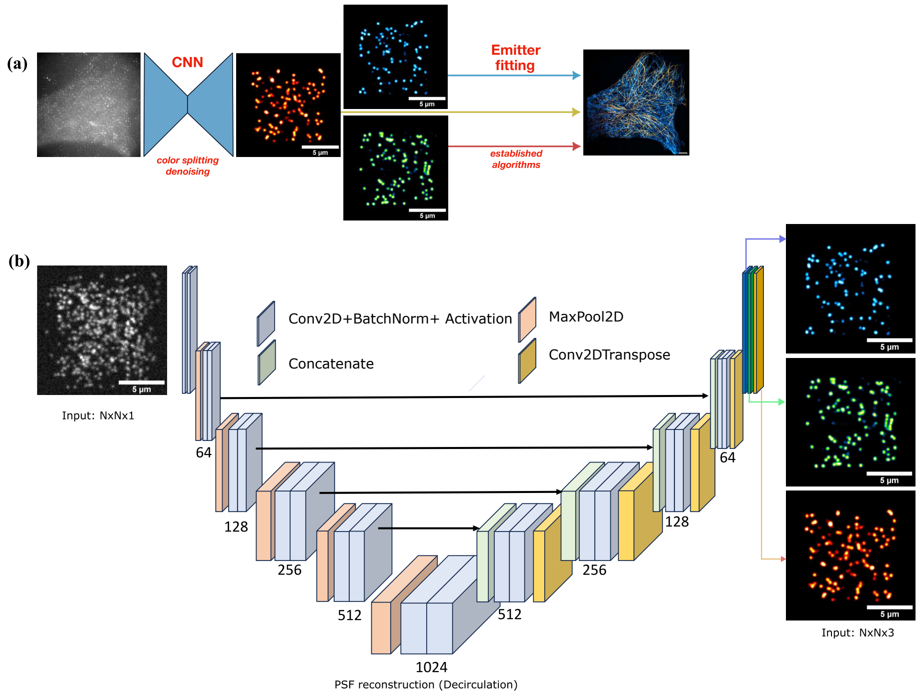

**Figure S6:** The outlined Decirculator approach. An input circulator image is analyzed with a U-net, trained to disentangle individual color channels and reconstruct the corresponding single lobe PSF. The network architecture is a slightly modified U-net architecture as described in Ref. [7].

to ELU (exponential linear unit) instead of ReLU (rectified linear unit) due to the depth of the network and to aid gradient propagation during training.

The model was trained on simulated data calculated to resemble the experimental data in terms of emitter brightness, PSF appearance, and background signal. We adapted a simulation pipeline that we described previously [8]. Briefly, we generated series of simulated images in which each image contained the fluorescence of a different set of randomly-positioned fluorophores of random emission colors (green, orange, or red). The number of fluorophores was furthermore chosen to meet a prescribed average emitter density. The set of allowed emitter densities was limited to 0.05, 0.1, 0.2, 0.4, 0.8, 1.5, 2.5, and 3.5 $\mu m^{-1}$, with a random density from this list chosen for each image. Each PSF lobe was modeled as a Gaussian function with standard deviations of 151 nm, 134 nm, and 115 nm for red, orange, and green, respectively, where each detector pixel integrated the corresponding part of this Gaussian over its active area. Based on our analysis of the experimental data, the total number of photons detected for each fluorophore was randomly sampled from a lognormal distribution function of the form

$$p(n, m, s) = \frac{1}{ms\sqrt{2\pi}} \exp\left\{ -\frac{[\ln(x) - m]^2}{2s^2} \right\} \tag{22}$$

with $(m, s)$ equal to (6.7, 0.27) (red), (6.9, 0.36) (orange), (6.65, 0.42) (green). After determining the expected number of photons detected in each pixel, we incorporated shot noise by replacing the pixel values with a random number drawn from a Poissonian distribution with a mean equal to the expected number of photons in that pixel. We then multipled this number by a factor 2.6, reflecting the conversion factor between photons and photoelectrons as provided by the camera manufacturer, and added a random number drawn from a normal probability distribution with a mean of 5000, reflecting the artificial and uniform offset of 5000 counts that was added in order to avoid the presence of negative values in the filtered data, and a standard deviation equal to 17.2329 to describe background.

Four different images were generated as training data: a simulated Circulator image in which the emitters appeared with the double-lobe PSF shape corresponding to their randomly-assigned emission colors, and three non-Circulator images in which the emitters of the corresponding color appear with the corresponding conventional (non-Circulator) PSF. The non-Circulator images did not include Poisson noise or background photons, allowing the CNN to also act as a denoising filter.

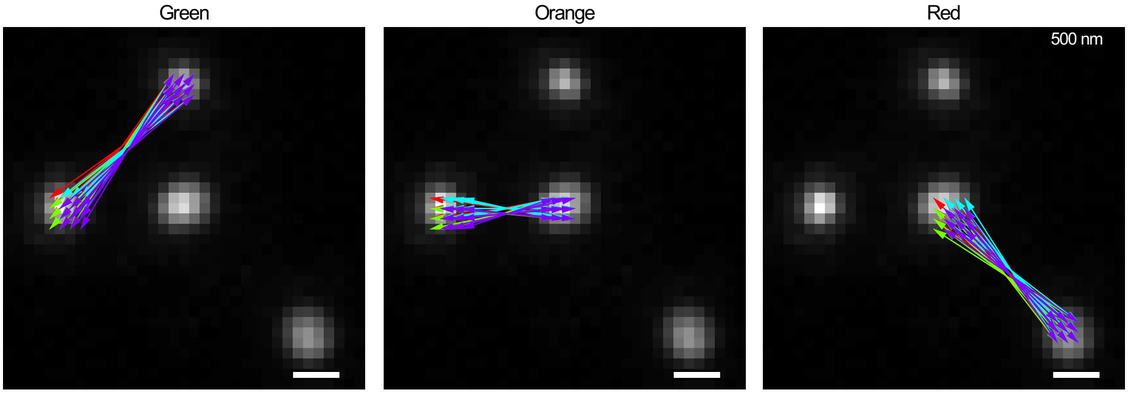

**Figure S7:** Cross-cumulant pixel combinations used to create SOFI images containing only the contributions of the green, orange, or red fluorophores.

The training of the CNN was performed by minimizing the following loss function:

$$L = \sum_i (y_i - \hat{y}_i)^2 + \alpha|\hat{y}_i| \tag{23}$$

where $y_i$ are the pixel intensities in the ground-truth non-Circulator images and $\hat{y}_i$ are the corresponding intensities predicted by the FCN. $\alpha$ is a regularizing parameter designed to promote sparsity, which was set to a value of $10^{-5}$. The total number of images for the training was set to 600,000 for each of the FCN's, with a 99/1% train-validation split respectively. Our network was trained for 1000 epochs and validation losses and weights were recorded at each epoch. The weights at the epoch with the lowest validation loss were taken for the final evaluation of the model performance. Once the validation loss started to increase, the training was halted as to avoid potential overfitting of the model.

The full SMLM analysis is then performed by applying a conventional SMLM analysis algorithm to all three of the CNN-generated images, providing the coordinates of the three different types of emitters.

Compared to the SMLM data, the live cell SPT data on IgE molecules was characterized by lower fluorescence brightnesses as well as the presence of molecules that were partially out of focus due to the deeper penetration depth required to visualize the apical plasma membrane. Accordingly, we decided to retrain the network on simulated data that better reflected the nature of the IgE data. In particular, we set the brightness of each type of fluorophore (green, orange, red) to half of the values used in training the SMLM data. To match the appearance of occasional brighter spots seen in the data, we also added a 15% chance that an emitter would be 1.7 times brighter than this baseline value. To model local defocus due to the non-flat nature of the cell membrane, we also increased the standard deviation of the PSF lobes by a factor 1.3, and scaled these values on a per-emitter basis with a random number sampled from a lognormal distribution function with $(m, s)$ equal to (0.36, 0.25). Finally, we added a background contribution similar to that observed in the data.

#### 4.3 SOFI analysis

SOFI imaging operates by calculating cross-cumulants of the intensities observed during the acquisition of multiple consecutive fluorescence images. A high-resolution SOFI image is then generated by creating an image in which the pixel values are those of the calculated cross-cumulants [9].

The cross-cumulants are closely related to the more common approach of cross-correlation, which can be easier to reason about intuitively. Cross-cumulants calculated for image series containing the fluctuating emission of blinking fluorophores will only be non-zero if the same fluorophore emits in all of the pixels involved in the cross-cumulant calculation. If the fluorophores emit into a set of distinct multilobe PSFs, it is therefore possible to define a set of cross-cumulants that are sensitive only to fluorophores emitting with one particular PSF shape, as was previously demonstrated for a 'double helix' PSF shape [10]. Figure S7 shows a graphical rendering of the cross-cumulants used to distinguish the different emission colors for the Circulator-created PSFs. The actual images were then calculated using our 'Localizer' software package [11].

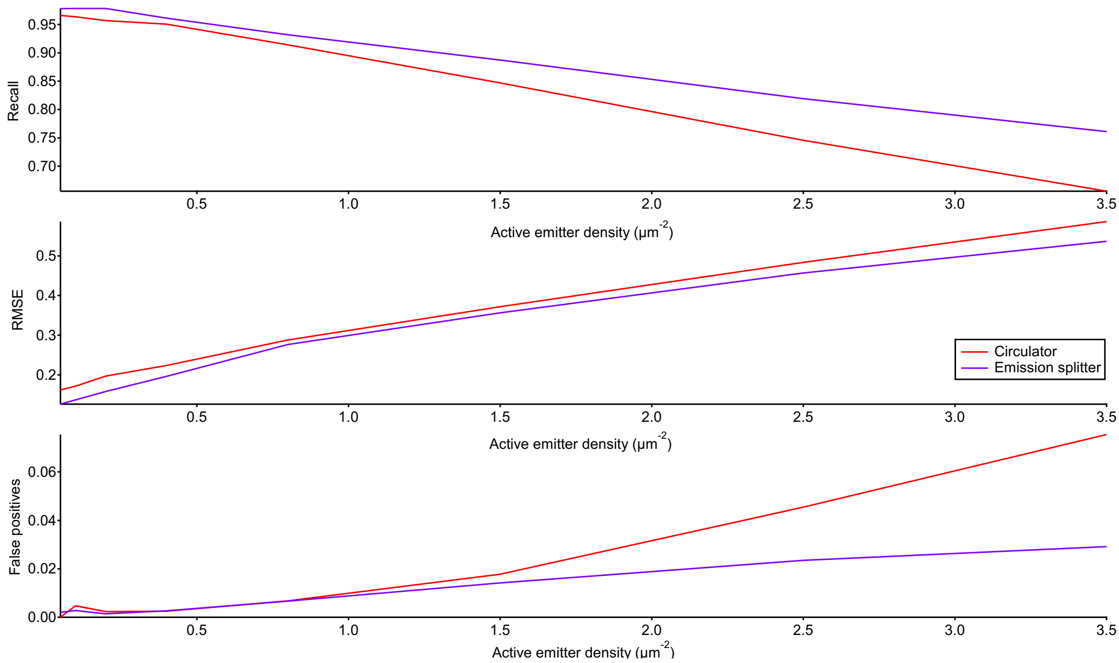

**Figure S8:** Simulated Circulator and image splitter SMLM performance for different emitter densities.

### Supplementary Note 5: Effect of the emitter density on the Circulator localization performance

The implementation of the Circulator described here doubles the number of emission spots that can be observed in a single image, which can make it difficult to discern individual emission spots. However, the degree to which this reduces the SMLM performance is not trivially clear because the Circulator PSFs are exactly known, and as a result there is latent information in the image that can be used to resolve individual emitters even from dense regions. We reasoned that this information may well be accessible to analysis methods such as the machine learning-based strategy detailed in Supplementary Note 4.2.

To evaluate the effect of the additional emission spots on the analysis accuracy, we set up a series of simulations designed to approximate measured fluorescence images, using the pipeline described in Supplementary Note 4.2. To provide a fair comparison, we also trained a CNN, entirely similar in architecture to the Circulator-analysis network described above, with the difference that it produced only a single output image. This network was trained to predict noise-free fluorescence images given data acquired without the Circulator. In this way the actions of this network mimicked the denoising inherent in the Circulator analysis CNN described in Supplementary Note 4.2, but for fluorescence data acquired using a single-lobe PSF.

We next generated test data in an entirely analogous way to that described in section 4.2, generating for each set of fluorophore coordinates a simulated fluorescence image using the Circulator and three different images containing the emission of only one color of fluorophores and unmodified PSFs.

We then applied the CNN-based analysis method described above to both analysis types. For the Circulator data, this meant the application of the pipeline described before, while for the non-Circulator data we applied the CNN that only implemented a denoising step followed by single-molecule localization using the same settings. The results of this analysis for a range of different emitter densities is shown in Figure S8. In this figure, we show three different performance parameters: the recall, which is the probability that a simulated fluorophore was correctly identified in the analysis, the root-mean square error of the localization estimate (RMSE), which shows how accurately the emitter position is determined, and the false positives fraction, which shows the fraction of all emitter localizations that are spurious, meaning that the analysis reported a fluorophore at a location where no simulated fluorophore was present. We judged that a localized emitter was correctly identified if there was a corresponding simulated fluorophore position within 200 nm.

As Figure S8 shows, the image splitting-based approach offers the highest performance across all three of the assessed parameters, though the Circulator offers a close performance, except for an increased propen-

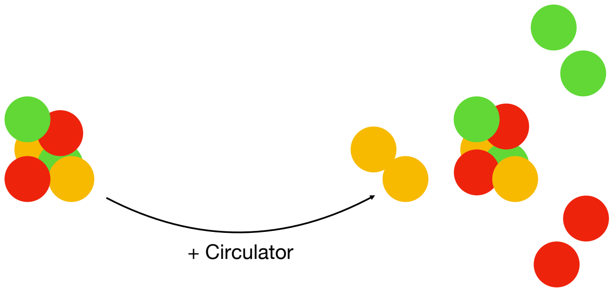

**Figure S9:** Conceptual illustration of how the Circulator spot duplication can help distinguish spatially-overlapping emitters due to the known structure present in the PSFs. The spots on the left would be difficult to untangle clearly, though this becomes easier when the Circulator PSF is used.

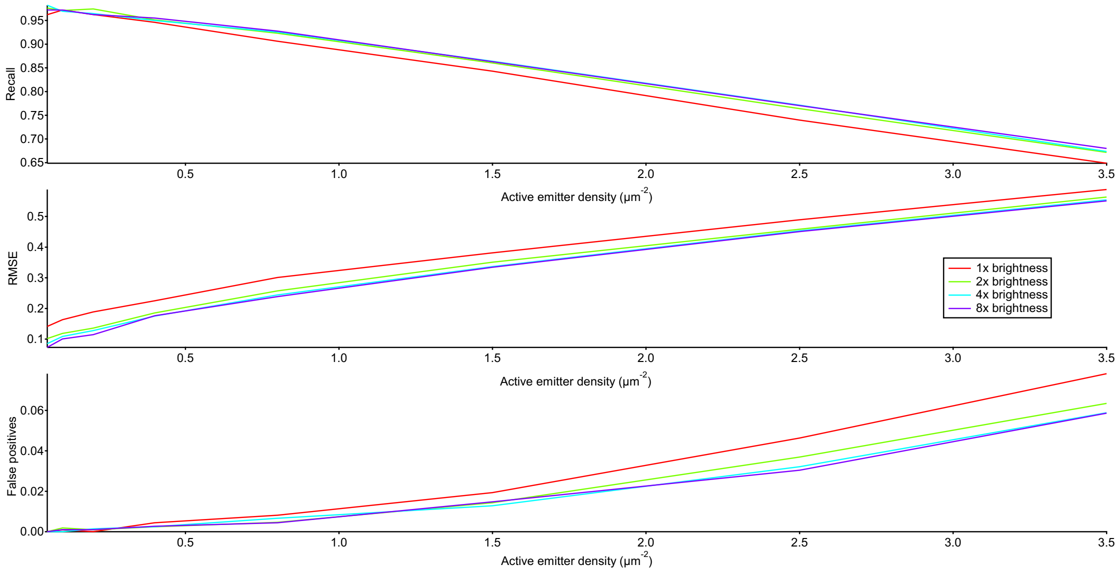

**Figure S10:** Circulator analysis performance using the CNN-based strategy for different emitter brightnesses and densities. A '1x' brightness is comparable to the brightnesses observed in our experimental data.

sity towards the generation of false positives though still within reasonable limits. The fact that the Circulator approximates the image-splitter performance may appear surprising at first sight. We interpret this as originating from the additional information present in the precisely-known PSF shapes, which results in additional possibilities to unravel emitter overlap and mitigates the increased number of emission spots. We illustrate this conceptually in Figure S9. The CNN can use this to disambiguate the raw fluorescence image and produce three different synthetic fluorescence images containing an (apparent) threefold lower emitter density that are more easily analyzed by conventional SMLM software.

The brightness of the fluorophores used here and in our actual data is low due to the limited performance of the green dye. We also investigated to what extent the SMLM performance improved if the brightness of the fluorophores is increased by a factor 2, 4, or 8. As Figure S10 shows, this does indeed lead to improved performance over the full range of emitter densities.

### Supplementary Note 6: Calculation of the IgE aggregate sizes

The number of IgE molecules in a complex (dimer or oligomer) and the average number of monomers in these complexes was calculated by considering the numbers of multicolor trajectories $m$, the number of single-color trajectories $s$, as well as the PSF patterns observed in the multicolor trajectories. We assumed that the green-, orange-, and red-labeled IgE molecules were present in similar concentrations on the sample, that no unlabeled molecules were present, and that the molecular association was not affected by the color of the labeling. In such a case, the color distribution of complexes consisting of $N$ labeled monomers is determined by multinomial statistics with equal probabilities for the occurrence of each color. We found that the ratio $r$ of the number of particles showing four emission lobes (containing at least one green and red fluorophore) to the number of particles showing three emission lobes (containing at least one orange fluorophore and at least one green or red fluorophore but not both) was characteristic for the average size of the associations, being given by

$$r = \frac{1 - 2(2/3)^N + (1/3)^N}{2(2/3)^N - 4(1/3)^N} \tag{24}$$

In particular, for dimers $r = 0.5$, for trimers $r = 1.0$, four-mers $r = 1.79$, and so forth based on simple enumeration of all possible combinations of labeled fluorophores.

This equation can be derived as follows. For a complex containing $N$ labeled IgE's, the probability of observing a single-color PSF is given by

$$p(n = 2) = 3\left(\frac{1}{3}\right)^N \tag{25}$$

since there are three possible colors. Let $n$ denote the number of lobes seen for a particular emitter. For multicolor particles, the probability of seeing three or four emission lobes in the PSF is given by

$$p(n = 4) = 1 - p(\text{no green}) - p(\text{no red}) + p(\text{no red and no green}) \tag{26}$$

$$= 1 - 2\left(\frac{2}{3}\right)^N + \left(\frac{1}{3}\right)^N \tag{27}$$

$$p(n = 3) = 1 - p(n = 2) - p(n = 4) \tag{28}$$

$$= 2\left(\frac{2}{3}\right)^N - 4\left(\frac{1}{3}\right)^N \tag{29}$$

Equation (24) is then simply given by the ratio of equations (27) and (29).

Once the average value of $N$ is known, the fraction of IgE molecules present in a complex $f$ was calculated using

$$f = \frac{mN}{s\,(1 - 3(1/3)^N) + m\,(N - 3(1/3)^N)} \tag{30}$$

which takes into account the fact that some multi-emitter particles will be classified as single-emitter particles if they result from the association of IgEs labeled with the same color. In the specific case of dimeric complexes found here, this equation reduces to

$$f = \frac{3m}{2.5m + s} \tag{31}$$

Equation (30) is obtained by considering that

$$f = \frac{mN/[1 - p(n = 2)]}{s + mN/[1 - p(n = 2)] - mp(n = 2)/[1 - p(n = 2)]} \tag{32}$$

where the factor $m/[1-p(n = 2)]$ corrects for the fact that some complexes appear as single-color emitters if they are labeled with dyes of the same color.

The fraction of all particles categorised as single color while having multiple same-color labels $g$ is given by

$$g = \frac{mp(n = 2)}{(m + s)[1 - p(n = 2)]} = \frac{m(1/3)^N}{(s + m)[1/3 - (1/3)^N]} \tag{33}$$

In the specific case of dimeric complexes found here, this equation reduces to

$$g = \frac{m}{2(m + s)} = \frac{f}{6 - 3f} \tag{34}$$

[7] Olaf Ronneberger, Philipp Fischer, and Thomas Brox. U-net: Convolutional networks for biomedical image segmentation. *CoRR*, abs/1505.04597, 2015.

[8] Wim Vandenberg and Peter Dedecker. Effect of probe diffusion on the SOFI imaging accuracy. *Sci Rep*, 7:44665, Mar 2017.

[9] Thomas Dertinger, Ryan Colyer, Robert Vogel, Jörg Enderlein, and Shimon Weiss. Achieving increased resolution and more pixels with superresolution optical fluctuation imaging (sofi). *Opt Express*, 18(18):18875–85, Aug 2010.

[10] A. Purohit, W. Vandenberg, T. Dertinger, D. Wöll, P. Dedecker, and J. Enderlein. Spatio-temporal correlation super-resolution optical fluctuation imaging. *Europhys. Lett.*, 125(2):20005, February 2019.

[11] Peter Dedecker, Sam Duwé, Robert K Neely, and Jin Zhang. Localizer: fast, accurate, open-source, and modular software package for superresolution microscopy. *J Biomed Opt*, 17(12):126008, Dec 2012.
